# A simple, open-source restraint system for magnetic resonance imaging in awake rats

**DOI:** 10.1101/2025.10.18.683153

**Authors:** Richard Quansah Amissah, Mahmoud Khaled Hanafy, Hakan Kayir, Peter Zeman, Kyle Gilbert, Alex Li, Miranda Bellyou, Ashley L. Schormans, Brian L. Allman, Jibran Khokhar

## Abstract

Magnetic resonance imaging (MRI) is a critical tool for translational neuroscience, offering cross-species insights into brain structure and function; however, its application in preclinical research is constrained by routine anesthesia use or sedation, which alters neural activity and limits comparisons to awake human imaging. Awake rodent functional MRI (fMRI) provides a powerful platform for investigating brain function under physiologically relevant conditions, but implementation is limited by technical challenges, particularly head motion and stress during scanning. Most restraint systems employ initial anesthesia, compromising translatability of findings, and highlighting the need for improved designs.

We developed a novel restraint system optimized for awake rat fMRI. The system consists of modular 3D-printed components and can be assembled in under five minutes. It is accompanied by a protocol that includes head-post implantation followed by an 11-day habituation period post-surgical recovery. The system eliminates the need for isoflurane anesthesia, ear bars, and bite bars, reducing stress and improving animal comfort. It supports integration with behavioral paradigms such as pupil tracking and licking responses.

High-resolution T2-weighted anatomical images and functional scans obtained using the system showed excellent spatial clarity and minimal motion artifacts. Quality control metrics, including head motion parameters and temporal signal-to-noise ratio, confirmed the system’s stability and suitability for awake imaging. Functional connectivity analysis revealed robust positive correlations between functionally relevant regions. This system offers a scalable, reproducible, and animal-friendly solution for awake rat fMRI. While the current design limits direct cranial access for multimodal recordings, it enables high-quality, behaviorally enriched imaging without anesthesia.

Significance Statement: Most rodent fMRI studies, including awake studies, rely on anesthesia, which profoundly alters brain activity and limits the interpretation of the data. This study presents a novel restraint system that enables high-quality fMRI in fully awake rats, eliminating the need for anesthesia, ear bars, and bite bars. By reducing stress and motion, this simple restraint system allows for investigation of neural activity and connectivity without confounds from sedation or anesthesia. Its open-source, modular design supports behavioral tasks and broad accessibility, making it a valuable tool for neuroscience research seeking to bridge the gap between preclinical imaging and real-world brain function.

## INTRODUCTION

Functional magnetic resonance imaging (fMRI) has become a valuable tool in translational neuroscience, offering non-invasive, whole-brain mapping of neural activity via blood-oxygen-level-dependent (BOLD) signal (Kwong et al., 1992). Unlike other methods such as electrophysiology, mini-scope imaging, and fiber photometry, fMRI provides superior spatial resolution and brain-wide coverage, enabling the investigation of distributed networks during rest and behavior, in addition to being akin to signals captured in human populations (Glover, 2011; Rizzolatti et al., 2018). Most rodent fMRI studies, however, rely on anesthesia or sedation to reduce motion, and stress during scanning. While effective for immobilization, anesthesia alters neurovascular coupling, suppresses neural activity, and disrupts functional connectivity, thereby limiting interpretability of functional imaging data (Sloan, 1998; Martin et al., 2006; Paasonen et al., 2018).

Awake rodent fMRI avoids these confounds, allowing researchers to examine brain function under more naturalistic conditions (Ferris, 2022). Despite its advantage, widespread adoption has been hindered by technical barriers, including procedural complexity, lack of standardized protocols, and reliance on custom hardware. Existing restraint systems for awake imaging are generally classified as invasive or non-invasive, each with trade-offs between motion control and animal welfare. While several systems have been developed for mice (Harris et al., 2015; Yoshida et al., 2016; Madularu et al., 2017; Chen et al., 2020; Fadel et al., 2022; Gutierrez-Barragan et al., 2022; Mikkelsen et al., 2022; Laxer et al., 2025), many require brief anesthesia during habituation and/or scanning. Restraint-based systems for rats remain relatively underdeveloped due to their larger size and greater strength, and often still depend on initial anesthesia (King et al., 2005; Febo, 2011; Upadhyay et al., 2011; Chang et al., 2016; Stenroos et al., 2018; Ma et al., 2020; Russo et al., 2021; Wallin et al., 2021).

In this study, we present a detailed, reproducible protocol for conducting awake fMRI in rats using a custom-designed, modular restraint system. Our approach combines skull-screw-less head-post implantation with a structured 11-day habituation regimen to minimize stress and motion without the use of anesthesia. The system is compatible with behavioral tasks and supports stable imaging suitable for longitudinal studies. By addressing both technical and welfare challenges, our method enhances the reproducibility and translational relevance of rodent fMRI research.

## MATERIALS AND METHODS

### Animals

All experimental procedures were approved by Western University’s Animal Care Committee (Approval Number: 2025-001) and were conducted in accordance with both provincial (Ontario Ministry of Agriculture, Food and Agribusiness; OMAFA) and federal (Canadian Council on Animal Care; CCAC) guidelines. Five approximately 6-month-old male (1 Sprague Dawley and 2 Long Evans) and female (2 Long Evans) rats were used for this study. Animals were housed in a controlled environment maintained at 21 ± 2 °C, with 30–40% relative humidity and a 12:12-hour light/dark cycle (lights on at 07:00). Standard rodent chow (14% protein, Envigo, WI, USA) and water were available ad libitum.

### Restraint System Design

The entire restraint system (Fig. 1) was 3D printed. With the exception of the head-fixing bar, which was printed using Tough 2000 Resin (FormLabs, Sommerville, MA, USA), all other components were printed with Tough Polylactic Acid (PLA; Shop3D, Mississauga, Canada). The head post and head post screw were printed using polyetheretherketone (PEEK) filament, while the plastic screws were printed from nylon filament (Shop3D, Mississauga, Canada). The head-fixing bar was printed over two days on a Form 3+ printer (FormLabs, Sommerville, MA, USA). The remaining PLA and polycarbonate components were printed on a RAISE3D Pro2 Plus printer (RAISE3D, Houston, Texas). After printing, the head-fixing bar was washed and cured. Brass screws were used to attach the connector to the cradle of the restraint system. The actual radiofrequency (RF) coil contains eight copper loops (Zanini et al., 2023). Post-processing of all parts included drilling and sanding. The entire restraint system was printed over the course of one week using the printers’ fastest settings. The total cost of the system components was approximately CAD $350.

**Figure 1.**
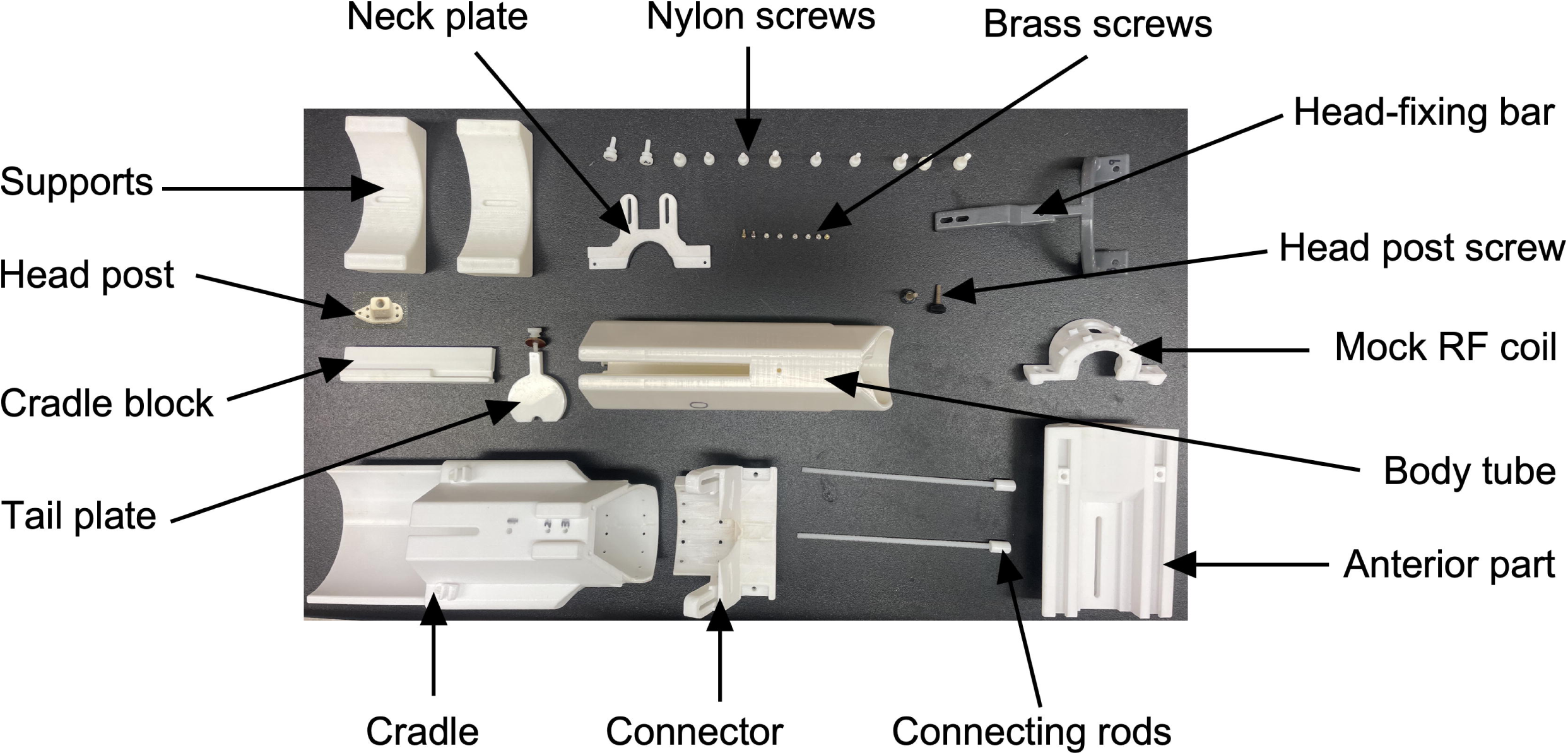
Components of the restraint system. The restraint system consists of 14 components: neck plate, nylon screws, brass screws, head-fixing bar, head post screw, mock radio frequency (RF) coil, body tube, anterior part, connecting rods, connector, cradle, tail plate, cradle block, and supports. The implanted head post used for head fixation is also shown. Except for the brass screws, all components were 3D-printed in-house.

Steps to assemble the restraint system (Fig. 2)

1. Place the cradle on a stable surface.
2. Attach the connector to the cradle using the brass screws.
3. Attach the anterior part of the restraint system to the cradle and connector using the connecting rods.
4. Insert the body tube into the cradle and secure it with a nylon screw.
5. Place the neck plate into the cavity between the cradle and connector, and fasten it with nylon screws.
6. Position the mock RF coil on the anterior part of the connector and attach it with nylon screws.
7. Place the tail plate in the groove on the body tube and tighten its screw.
8. Fix the head-fixing bar to the anterior part of the restraint system. Ensure that the head post receptacle is aligned directly above the hole in the mock RF coil.
9. Secure the head-fixing bar to the cradle with nylon screws.

**Figure 2.**
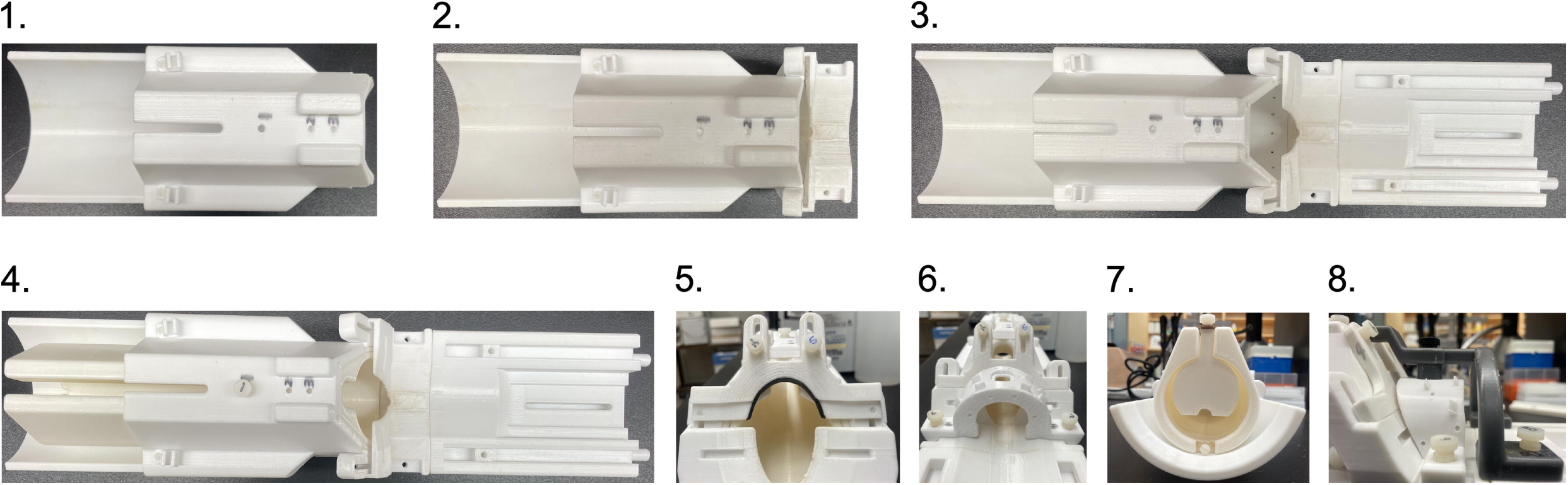
Assembly of the restraint system. Step-by-step images illustrating the simple, 8-step procedure for assembling the restraint system.

**Figure 3.**
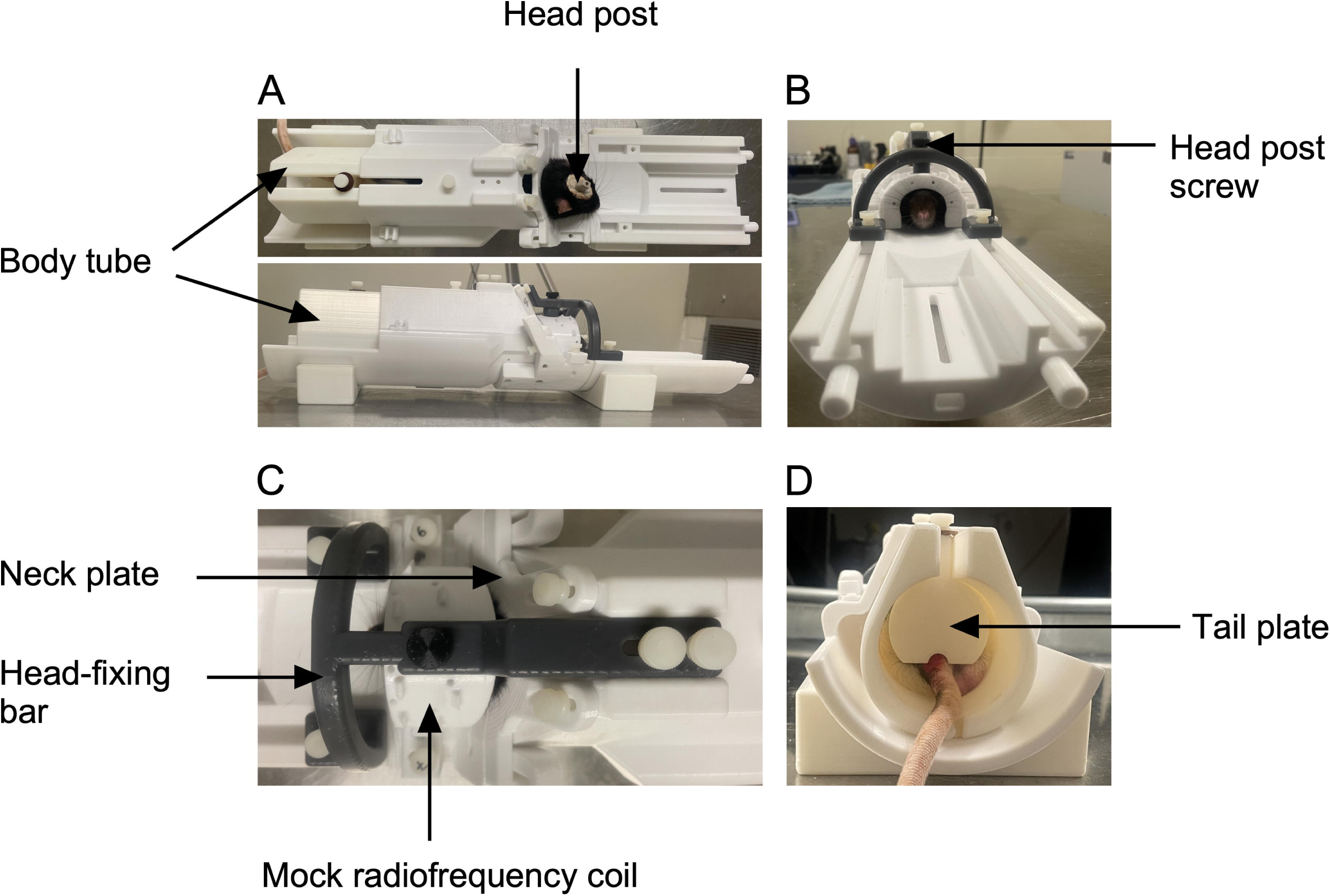
Head fixation of a rat in the restraint system (multiple views). A. Top panel: Rat positioned in the restraint system with the neck plate secured. Bottom panel: Side view of the system after placement of the head-fixing bar, which is attached to the rat via the implanted head post and secured to the restraint system. B. Front view showing the head-fixing bar secured to both the rat and the restraint system. C. Top view showing details of the nylon screws and head post screw used to firmly immobilize the rat’s head. D. Rear view illustrating the tail plate and positioning of the rat’s tail during head fixation and scanning.

### Stereotaxic Surgery

The rat was first weighed to calculate the appropriate dose of Metacam (meloxicam oral suspension, Boehringer Ingelheim Vetmedica GmbH) and Baytril (enrofloxacin, Elanco US Inc., Greenfield, IN, USA) for administration during surgery. Anesthesia was induced with 5% isoflurane and 2% oxygen until the desired depth of anesthesia was reached. The rat was then injected with meloxicam (5 mg/kg) and Baytril (5 mg/kg). Next, the rat’s head was shaved to expose the scalp. The rat was positioned on the stereotaxic apparatus, with a pre-warmed heating pad used to maintain body temperature. A rectal probe was inserted to monitor body temperature continuously. Ear bars and a bite bar were used to stabilize the head. Eye gel was applied to prevent corneal drying. The shaved skin was disinfected sequentially with soap, isopropyl, and chlorhexidine to maintain a sterile field.

The isoflurane concentration was then reduced to 2%. A toe pinch was performed to confirm the absence of reflexes and ensure a sufficient depth of anesthesia. A midline scalp incision was made from the anterior to the posterior aspect of the head to expose the skull. Underlying tissues were removed, and the skull was cleaned to clearly expose the bregma and lambda. Etchings were made on the skull to enhance bone cement adhesion. The head post, stored in isopropyl until needed, was mounted onto the vertical arm of the stereotaxic apparatus and positioned over the lambda. Care was taken to avoid tilting the head post to ensure proper orientation during MRI scanning. A thin layer of All-Bond Universal (BISCO Inc., Canada) was applied to the skull to improve bone cement adhesion and cured with UV light. Bone cement (Core-Flow DC Lite, BISCO Inc, IL, USA) was then applied. The head post was lowered onto the skull so that its lower portion was embedded within the cement. Later, the thickened, yet still fluid, bone cement was smoothed using a spatula to eliminate sharp edges that could cause irritation after hardening, and then cured.

Once the cement had hardened, the head post was detached from the stereotaxic arm. The scalp was repositioned over the bone cement, without requiring sutures. The area was cleaned of residual blood. Finally, 3 ml of saline was administered subcutaneously. Isoflurane was gradually turned off, while oxygen was maintained until the rat regained consciousness. The ear and bite bars were quickly removed at the first signs of regained consciousness. The rat was transferred to a recovery cage until it was fully mobile, and then placed in a clean cage. Post-operative care included administration of Metacam and Baytril for three days. The rat was allowed a 10-day recovery period before restraint system habituation began.

Steps for habituation to restraint, head fixing, and scanner noise

This protocol outlines a step-by-step method to habituate rats to restraint, head fixation, and scanner noise for awake fMRI studies. It is designed to minimize stress and movement while preserving physiologically relevant brain states. The entire process spans about 11 days.

Before beginning the restraint protocol, habituate the rat to handling, head-holding by hand, and receiving a reward after restraint and head-fixing. Gradually increase the duration of the head-holding until you can comfortably hold the rat’s head for about one minute.

1. Habituation to the restraint system (Days 1 & 2): Place the restraint system in an open field box and allow the rat to explore it for 10 minutes each day.
2. Body immobilization (Days 3 & 4): Immobilize the rat’s body using the body tube, leaving the head unrestrained.

i. Place the body tube halfway on a regular napkin, with the rear of the tube resting on the napkin.
ii. Position the rat on the napkin with its head facing the rear of the body tube.
iii. Wrap the napkin around the rear of the tube and the rat to encourage the rat to move forward through.
iv. Once the rat enters the tube, lift the tube and gently insert it into the cradle.
v. Raise the neck plate to encourage the rat to extend its head through the front opening of the cradle. If needed, use the tail plate to gently push the rat forward.
vi. Once the rat positions its head under the neck plate, gently lower the neckplate and secure it with screws to prevent backward movement.
vii. Insert the tail plate into its designated groove to prevent backward movement, and secure it tightly. Ensure the rat’s tail passes through the tail plate grove to avoid injury.
viii. Keep the rat in this position for 15 minutes on day 3 and 30 minutes on day 4.
3. Body immobilization with head fixing (Days 5 & 6): Immobilize the rat as previously described and head-fix it.

i. With the rat’s head protruding from the neck plate, place the mock RF coil over the head so that the head post extends through the hole in the coil. Secure the coil to the restraint system.
ii. Position the head-fixing bar so that its designated slot is directly above the head post.
iii. Secure the head post to the head-fixing bar to prevent both vertical and horizontal head movement.
iv. Attach the front and rear parts of the head-fixing bar to the restraint system to secure it.
v. Keep the rat immobilized and head-fixed for 15 minutes on day 5 and 30 minutes on day 6.
4. Immobilization with head-fixing in mock MRI chamber (Days 7 to 11): After immobilization and head fixing, place the rat with the restraint system inside the mock MRI chamber and expose it to recorded MRI sounds.

i. Using a speaker capable of producing sounds up to 120 dB sound pressure level (SPL; the intensity of sounds produced by the MRI during scanning), play the recorded MRI sounds–starting at a lower intensity on day 7 and gradually increasing the intensity each day, reaching full intensity by day 11.
ii. From days 7 to 11, immobilize the rat with head-fixing in the mock MRI chamber for progressively longer durations: 30, 45, 60, 90, and 120 minutes, respectively.

After completing the 120-minute restraint, the rat can be placed in the actual MRI scanner. However, repeating the two-hour restraint session several more times is recommended to further habituate the rat and reduce stress during actual scanning. Additionally, providing a reward after each restraint and/or head-fixing session is encouraged to create a positive association with the experience.

### Mock MRI chamber

The mock MRI chamber used in this study was designed as a double-walled sound booth, consisting of two MedAssociates (Burlington, VT, USA) sound-attenuating cabinets of different sizes, with the smaller, internal chamber (ENV-022MD; Standard MDF cubicle) placed inside the larger one (ENV-017; Extra-large MDF cubicle). The interior walls of the internal chamber were lined with soundproofing foam sheets (2” pyramidal shape; Foam Factory, Macomb, MI, USA) to limit sound reflections and to minimize sound escaping from the small portals in the mock MRI chamber used for cable passage. Inside the chamber, a speaker capable of producing sound levels up to 120 dB SPL was installed. The speaker was connected to an external laptop used to play the recorded MRI sound file. The MRI sounds were recorded from a 9.4T Bruker MRI scanner and included noises produced during shimming, T2-weighted image acquisition, and fMRI acquisition. The total duration of the recorded sounds was 2 hours. Depending on the stage of habituation, the appropriate segment of the recording was played.

### MRI Experiments

Imaging was performed at the Center for Functional and Metabolic Mapping, located within the Robarts Research Institute at Western University. Scans were obtained using a Bruker 9.4T, 31-cm horizontal-bore magnet (Varian/Agilent, Yarnton, UK) equipped with a 6-cm Magnex HD gradient insert and a Bruker BioSpec Avance NEO console running ParaVision 360 v3.3 (Bruker BioSpin Corp, Billerica, MA). T2-weighted anatomical images were acquired at the start of each session using a TurboRARE pulse sequence (8 averages, 35 slices, slice thickness = 400 μm, field of view (FOV) = 38.4 x 38.4 mm, matrix size = 192 x 192, in-plane resolution = 200 x 200 μm, repetition time (TR) = 1500 ms, echo time (TE) = 15 ms) (Hennig et al., 1986). Resting-state fMRI data were collected using a gradient-echo echo-planar imaging (EPI) sequence with optimized parameters (Gilbert et al., 2019): 4 runs of 400 volumes each, TR = 1500 ms, TE = 15 ms, FOV = 38.4 x 38.4 mm, matrix size = 96 x 96, 35 slices, isotropic resolution = 400 μm, bandwidth = 280 kHz). Image processing was performed using the Rodent Automated Bold Improvement of EPI Sequences (RABIES) software in conjunction with the SIGMA rat brain template (Barrière et al., 2019; Desrosiers-Grégoire et al., 2024). A detailed description of the workflow used for fMRI data processing was described in a previous study (Grandjean et al., 2023; Frie et al., 2024).

## RESULTS

Following the design and 3D printing of the rat restraint system, rats were habituated to restraint and head-fixing over a period of 11 days. After habituation, rats were placed in the MRI scanner for imaging. Each session began with T2-weighted image acquisition, followed by four fMRI runs. The full scanning session lasted up to two hours, during which the rat remained awake and motion-restricted. Representative T2-weighted and functional MRI images from an awake rat are shown in Figs. 4A and B, respectively.

**Figure 4.**
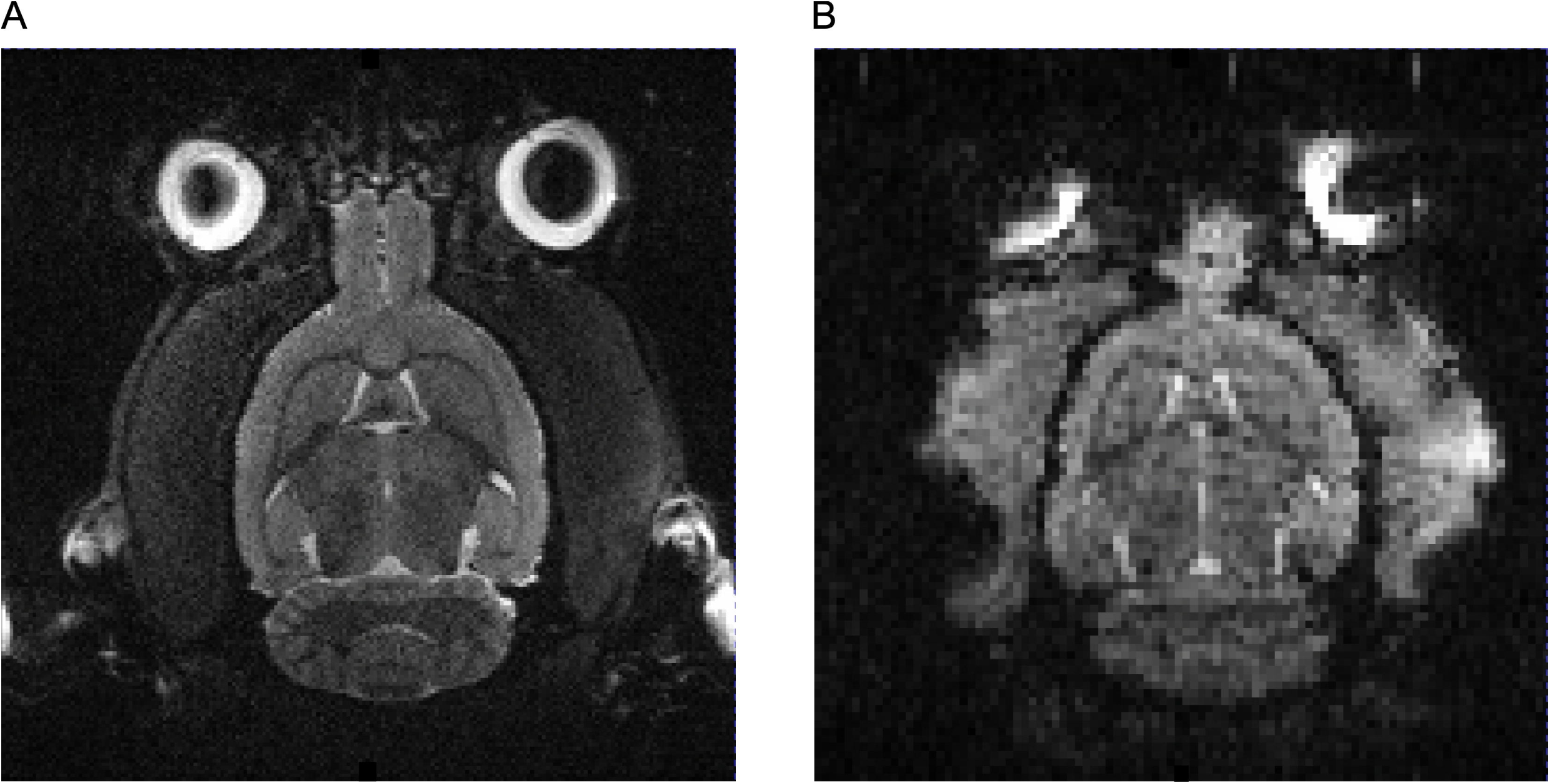
MRI images acquired using the rat restraint system. A. T2-weighted anatomical image B. Functional MRI image from the first volume of the first scanning run in an awake, head-fixed rat.

The T2-weighted axial image demonstrates high anatomical contrast and clear delineation of major brain structures, including the neocortex, cerebellum, and ventricular spaces, with minimal evidence of motion artifacts. The corresponding fMRI slice shows adequate signal homogeneity and preservation of anatomical landmarks despite lower spatial resolution, supporting reliable co-registration with the structural image. Minor susceptibility-related artifacts are visible near the eyes and olfactory bulbs, as expected with echo-planar imaging, but no significant motion-related distortions are observed. Together, these images confirm that the restraint system and habituation protocol enabled the acquisition of high-quality structural and functional data in fully awake rats without the use of anesthesia.

Head motion and signal quality metrics were assessed using the RABIES preprocessing pipeline. As shown in Figure 5, the subject exhibited minimal head motion throughout the scan. Translation parameters remained within ±0.05 mm, while rotational displacements were confined to ±0.005 radians, indicating stable positioning throughout the acquisition period. Framewise displacement values were consistently low, not exceeding 0.005 mm, with very few spikes, suggesting the absence of significant motion artifacts. Temporal standard deviation maps revealed the expected spatial distribution of signal variability, with higher values localized to cortical and subcortical regions. These areas typically exhibit greater BOLD fluctuations due to physiological activity. Temporal signal-to-noise ratio maps demonstrated robust signal quality across the brain, with values exceeding 30 in most regions. These results confirm that the data quality is sufficient for subsequent analyses, with minimal noise contamination and adequate BOLD sensitivity across the scanned volume.

**Figure 5.**
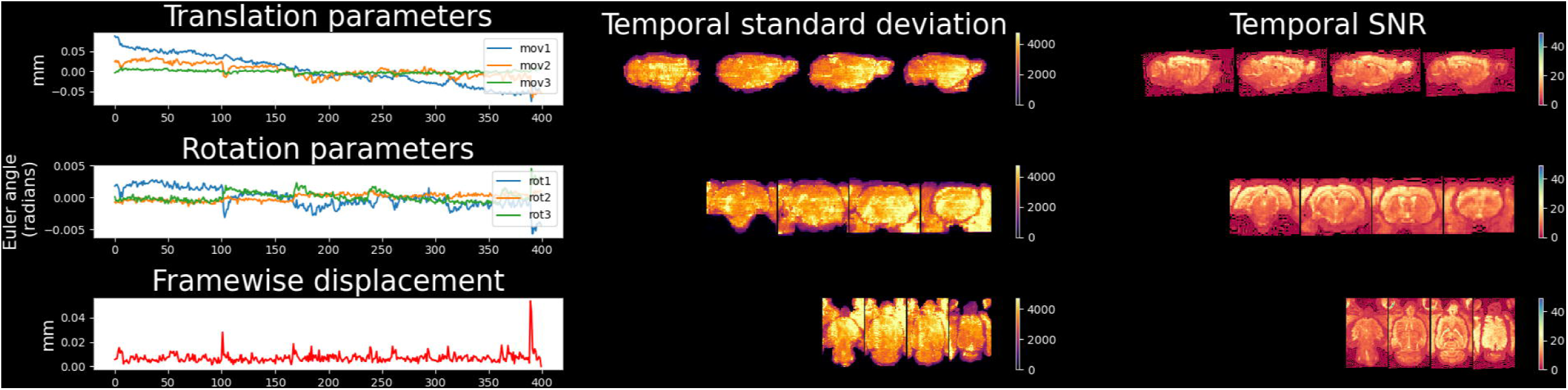
Head motion and image quality metrics during fMRI scanning. Left panel: Top: Time course of translational head motion parameters, Middle: Rotational motion parameters, and Bottom: Framewise displacement across the scanning session. Middle panel: Temporal standard deviation maps displayed in sagittal (top), coronal (middle), and axial (bottom) views. Right panel: Temporal signal-to-noise ratio maps for sagittal (top), coronal (middle), and axial (bottom) views.

**Figure 6.**
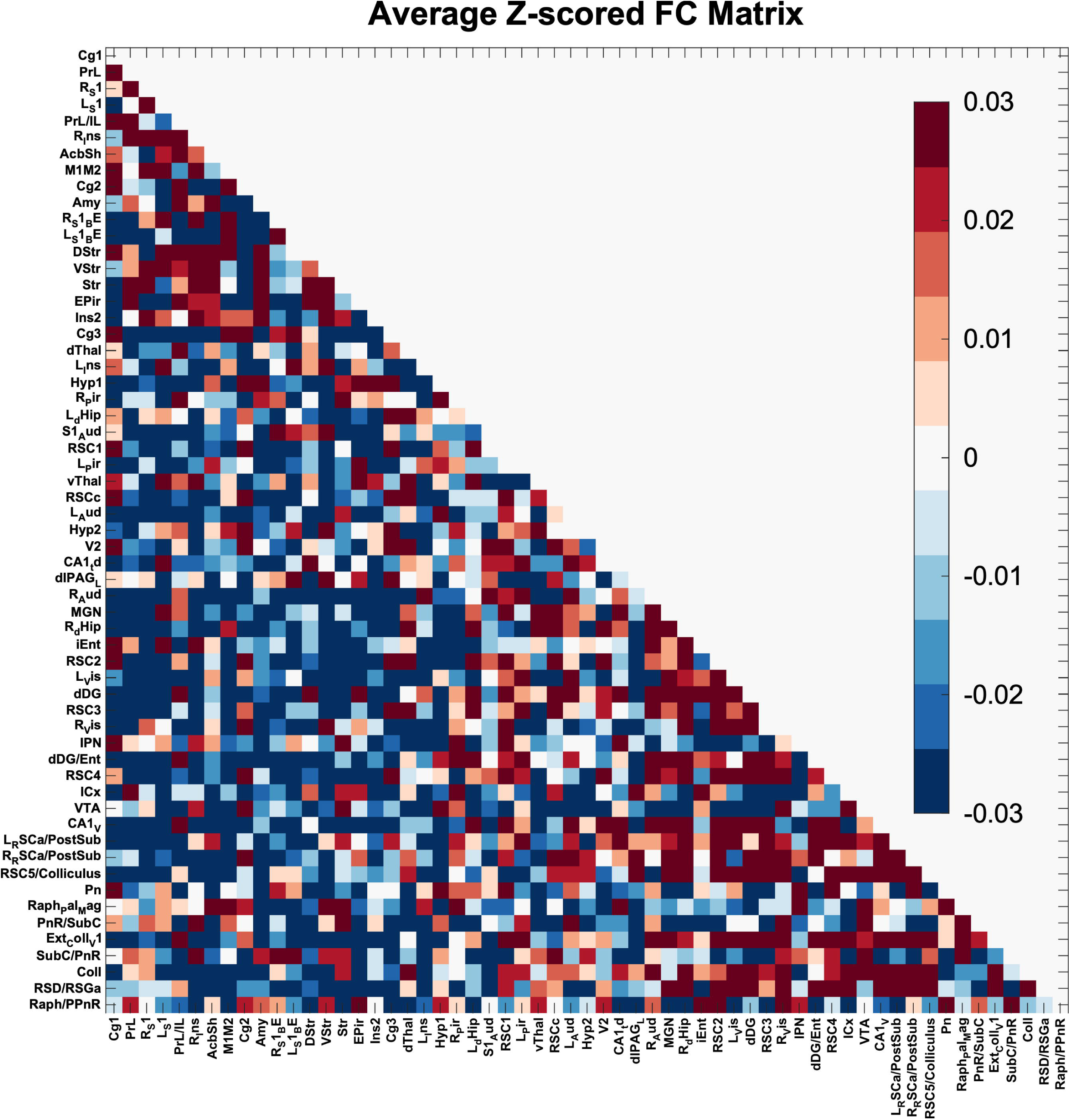
Average Z-scored functional connectivity matrix in awake rats. The matrix shows pairwise Z-transformed Pearson correlation coefficients between BOLD signals across 59 brain regions. Warmer colors indicate stronger positive connectivity, and cooler colors indicate weaker or negative connectivity.

We generated a Z-scored functional connectivity (FC) matrix from resting-state fMRI data in awake rats, revealing structured patterns of both positive and negative connectivity across brain regions. Positive correlations were evident among limbic and associative areas, including the prelimbic cortex (PrL), cingulate cortex (Cg1), retrosplenial cortex (RSC), hippocampus, and entorhinal cortex—consistent with a default-mode-like network (Upadhyay et al., 2011). In contrast, subcortical and brainstem regions such as the thalamus, periaqueductal gray (PAG), and Raphe nuclei exhibited weak or negative connectivity with cortical areas, suggesting functional segregation during rest. The overall range of FC values was modest (±0.03 Z-scores), indicating subtle but topographically organized network architecture in the awake rat brain.

## DISCUSSION

Our rat restraint system was designed to greatly minimize head motion and ensure animal comfort, without induction anesthesia use, thereby reducing stress to enable the acquisition of high-quality images. The entire system was 3D-printed over the course of one week, is easy to assemble, and can be modified to support additional behavioral tasks during scanning. The habituation protocol is straightforward and reliably acclimates rats to awake scanning within 11 days.

Prior to restraint system use, surgery is required to implant a head post. Although invasive, this approach remains the gold standard for studies requiring the highest spatial and temporal resolution. Non-invasive systems may be preferred as they eliminate the need for surgery (Stenroos et al., 2018; Ferris, 2022); however, they typically lack the mechanical rigidity needed to prevent submillimeter head motion, particularly during extended scans or behavioral task engagement. In contrast, head-post-based systems provide superior head immobilization, facilitating the acquisition of motion-free images across long sessions and enabling precise image registration across time points (King et al., 2005; Tsurugizawa et al., 2020). Despite the surgical requirements, invasive systems remain the most reliable option when data fidelity, spatial accuracy, and compatibility with multimodal techniques are essential. The performance of our system demonstrates this advantage, yielding high-resolution T2-weighted and functional images with minimal motion artifacts.

The functional connectivity patterns observed in awake rats are consistent with previous reports describing intrinsic brain organization and network segregation in the rodent brain. The presence of positive connectivity among regions including the prelimbic cortex, cingulate cortex, and hippocampus, supports the existence of a default-mode-like network, which has been identified in awake rodents using both seed-based and independent component analyses (Upadhyay et al., 2011; Liang et al., 2012). This network is thought to underlie internally-oriented cognitive processes and is analogous to the human default mode network (Greicius et al., 2003), reinforcing the translational advantage of rodent fMRI. Conversely, the relatively weak or negative connectivity between cortical and subcortical regions, including the thalamus, periaqueductal gray, colliculi, and Raphe nuclei, suggests functional segregation of arousal- and motor-related systems from higher-order associative networks during rest. These findings align with prior work showing that awake-state functional connectivity is more modular and selective than under anesthesia, where widespread global synchrony often dominates (Paasonen et al., 2018). Together, the observed network topology highlights the capacity of awake rodent fMRI to capture biologically meaningful patterns of brain organization and provides a valuable reference point for evaluating circuit-level alterations in experimental models.

The development of effective restraint systems is central to advancing awake rodent fMRI, given the need to balance high-quality imaging with minimal stress and maximal animal welfare. Our system addresses several limitations inherent in existing designs by eliminating anesthesia use, reducing animal discomfort, and enabling behavioral integration during imaging. The avoidance of isoflurane is particularly critical, as even low doses have been shown to alter neural activity and functional connectivity in rodents, potentially confounding results in studies aimed at capturing brain dynamics in naturalistic, awake conditions due to the remaining effects of induction anesthesia (Sloan, 1998; Martin et al., 2006; Paasonen et al., 2018). Instead, our approach supports scanning under fully conscious conditions, enabled by a restraint design that reduces head and body motion during imaging sessions.

Unlike traditional systems that rely on ear bars and bite bars for head immobilization, our design eliminates these features entirely. This modification significantly reduces discomfort, a known source of stress and motion in head-fixed preparations. The system’s modularity and 3D-printed components allow for scalable adaptation, such as interchangeable tube inserts that accommodate rats of various sizes, sex, and ages, facilitating longitudinal studies and developmental imaging protocols. Another key advantage is the system’s compatibility with behavioral paradigms: integrated features support active tasks such as treadmill walking, which can serve as an operandum, similar to that used in a previous study (Vasilev et al., 2022). Because most of the rat’s face, including the eyes, remains accessible during restraint, additional behaviors such as pupil tracking, licking, and whisker movement can also be measured during scanning. This enables simultaneous acquisition of brain-wide activity and behavior in awake, task-engaged animals (Lawen et al., 2025), and integration with other open-source tools like fiber optic lickometers (Silva et al., 2024), or ability to measure the same consummatory behaviours in home-cage and in-magnet (Frie and Khokhar, 2024). The streamlined fixation process also contributes to experimental efficiency, typically requiring only about five minutes to prepare and position each rat for scanning.

Despite these strengths, the restraint system is not without limitations. Most notably, it restricts direct access to the skull, thereby limiting compatibility with techniques such as electrophysiology, optogenetics, or fiber photometry that require chronic implants or optical access. This constraint could be addressed by designing head posts that anchor around the periphery of the skull while leaving the central area accessible for multimodal data acquisition, or incorporating non-ferromagnetic inserts into the head-post design (Quansah Amissah et al., 2023). Additionally, although the restraint minimizes gross head motion, fine-scale movements such as whisking and jaw motion can still introduce artifacts. While these were infrequent and manageable in our experience, further reduction may be possible through additional behavioral training. Acoustic noise during MRI scanning remains another concern, as it can be a stressor for awake animals. To mitigate this, we recommend noise attenuation strategies such as silicone earplugs. One of the primary sources of MRI-related acoustic noise is the rapid switching of magnetic field gradients (MacKinnon et al., 2025). A promising technique to address this is Steady-State-On-the-Ramp Detection of Induction Decay with Oversampling (SORDINO), which maintains a constant gradient throughout scanning (MacKinnon et al., 2025). In addition to significantly reducing acoustic noise, SORDINO is resistant to motion and susceptibility artifacts, thereby improving image quality by minimizing both stress-induced movement and signal distortions.

Our restraint system represents a significant step toward behaviorally compatible and reproducible awake rodent fMRI, especially given the lack of anesthesia use. By improving comfort, reducing motion, and supporting a range of behavioral tasks, it enables studies that closely approximate natural brain states. Future iterations may seek to enhance access for multimodal applications (e.g., electrophysiology; optical imaging) and further mitigate the impact of residual motion and acoustic stressors.

## Acknowledgements

We are grateful to Amr Eed for sharing the magnet audio as well as post implantation methods. We would also like to acknowledge the support from the BrainsCAN imaging core funded by the CFREF-APOGEE.

